# Parametric response mapping as a new MRI approach for the characterization of post-traumatic cerebral edema

**DOI:** 10.1101/492298

**Authors:** Jules Grèze, Pierre Bouzat, Jean-François Payen, Emmanuel Barbier, Benjamin Lemasson

## Abstract

**Purpose:** Cerebral edema is usually assessed with mean ADC value obtained with MRI. We aimed to use parametric response mapping (PRM), a voxel-based analysis, in a rat model of traumatic brain injury (TBI) to characterize the cerebral edema and to compare findings to those obtained with the classical global mean ADC analysis. Ultimately we wished to test whether early PRM analysis could identify the vasogenic or cellular edematous pattern two hours after trauma.

**Materials and Methods:** Experiments were conducted in a rat model of TBI in accordance with French Government guidelines. Eighteen traumatic brain-injured rats (TBI group) were compared to 7 sham-operated rats (Sham group). Diffusion-weighted images were acquired before and immediately (H0), 60min (H1) and 120min (H2) after the trauma. Two regions of interest (ROI) including cortex (cortical-ROI) and the whole brain (brain-ROI) were manually delineated.

**Results:** Averaged ADC value in cortical-ROI was significantly reduced at H2 in the TBI group versus Sham group (715±24 vs 781±16 μm^2^/sec; p=0.04), not in brain-ROI (748±27 vs 801±18 μm^2^/sec; p=0.32). By contrast, PRM values were significantly changed in these two ROIs at as soon as H1 in the TBI group (27.5±4% of the voxels beyond the PRM threshold in TBI group vs 5.1±1% in sham group; p<0.01). PRM was able to identify the nature of post traumatic cerebral edema: vasogenic, cellular or mixed pattern. ADC data obtained immediately after TBI and analyzed using PRM correlated with ADC data obtained at H2.

**Conclusion:** The PRM approach is more sensitive than averaged ADC at detecting and identifying post-traumatic brain edema. It is able to PRM might be a promising tool to individualize the management of patients with posttraumatic brain edema.

## Introduction

Traumatic brain injury (TBI) is a major public health care issue, accounting for 235,000 hospitalizations per year in the USA (1). Development of brain edema following TBI is one of the determinants for secondary brain injuries that add to the burden of initial lesions by increasing intra-cranial pressure (ICP) and compromising cerebral hemodynamics. Two types of cerebral edema have been described since 1967: the vasogenic edema, relating to blood-brain-barrier leakage, and the cellular edema, induced by the accumulation of water and electrolytes inside the cell (2).

Monitoring post-traumatic cerebral edema remains challenging. The apparent diffusion coefficient (ADC) of water, measured with diffusion-weighted magnetic resonance imaging (MRI), is a valid marker to characterize cerebral edema. This parameter may be used to distinguish between vasogenic edema, as reflected by increased ADC values in the cerebral tissue, and cellular edema with decreased ADC values (3–5). In preclinical studies, ADC values are usually averaged across regions of interest (ROI), which are classically restricted to the cortical area, to obtain information about the type of cerebral edema (3, 6, 7). However, this approach does not account for the possible heterogeneity of cerebral edema within the brain tissue (8)).

In experimental models of TBI, different patterns of ADC over time have been observed during the development of post-traumatic brain edema, and the results have not been univocal. Focal TBI models with cortical cryogenic lesion induced vasogenic edema whereas controlled cortical impact models obtained prominent cellular edema within the core lesion and vasogenic edema in the surrounding area (9,10)). Following diffuse TBI induced by a weight-drop model, one study reported a bi-modal evolution of ADC values with an initial increasing of ADC followed by its decrease (3). Other authors, however, found a global decrease of ADC in this same model (5). This heterogeneity has been well described by Long et al. (11) as their observed in the hyper-acute phase (3 h) of TBI, the heterogeneous ADC decreases and increases around the impacted area indicated a combination of cytotoxic and vasogenic edema, respectively. To go beyond a this observation Li et al. proposed an original approach to quantify this heterogeneity(12)). They used several ROIs drawn within the cortex to quantify the heterogeneity of the edema at one given time. Despite the fact that this study described more precisely the brain response to a TBI this technique of quantification does not characterize the evolution of this heterogeneity overtime. Collectively, these results suggest that measuring a uniform ADC value within a ROI, and following this ROI overtime dilute the heterogeneous nature of the edema that occurs in different TBI models.

The heterogeneity of ADC values within brain pathology and its evolution overtime has recently been addressed in tumor studies. In one such study, a functional diffusion map approach was used to evaluate the ADC modifications over time on a voxel-wise basis in brain cancer patients (10). Values of ADC measured using this voxel-based approach were associated with the subsequent evolution of the tumor, i.e., early changes in tumor ADC values could be used as a prognostic indicator of subsequent volumetric tumor response, whereas averaged ADC values in brain tissue had no predictive value (11). Galbán et al. generalized this voxel-based quantification to perfusion MRI and successfully predicted the survival of patients with brain cancer (12). This voxel-based image analysis technique, known as parametric response mapping (PRM), could provide a more accurate means to assess the heterogeneity of cerebral edema, than using averaged ADC values after brain insult. Our intent was to implement the PRM approach in a well known rodent model of diffuse TBI (13, 14) to assess its ability to differentiate the two components of the post-traumatic cerebral edema. The primary aim of this study was to characterize cerebral edema and its evolution in a rat model of traumatic brain injury using both the PRM approach and the averaged ADC analysis.

## Material and Methods

### Experimental Protocol

The study design received approval from the local Internal Evaluation Committee for Animal Welfare and Rights. Experiments were conducted in accordance with French Government guidelines (licenses 380819 and A3851610008). A series of experiments with two groups of adult male Wistar rats (350-450g) was conducted, including a TBI group (n = 18 rats) and a Sham-operated group (n = 7 rats). Anesthesia was induced and maintained with isoflurane (inhaled fraction of 2.0%) during the study period. After tracheal intubation, rats were mechanically ventilated with 60% air–40% oxygen using a rodent ventilator (SAR-830/P, CWE, Ardmore, PA). Catheters were inserted into the femoral artery and vein for blood gases (PaO_2_, PaCO_2_) and blood pressure monitoring, and for the administration of fluids. Rectal temperature was kept at 36.5±0.5°C using a heating device. As per the initial description of impact-acceleration model (13, 14), TBI was induced by dropping a 500g mass through a vertical cylinder from a height of 1.5m onto a metallic disc glued to the skull. The reference time (H.ref) corresponded to the first MRI acquired before trauma (or equivalent time) in the two groups of rats.

### MRI Measurements

MRI was performed at 7T in a horizontal bore magnet (Bruker Biospec 47/40 USR AV III, Bruker BioSpin, France; IRMaGe facility, Grenoble, France) equipped with actively shielded gradient coils generating gradients of up to 600mT/m. Four anatomical (T2w) and diffusion-weighted images (DWI) were acquired in the TBI group: before the trauma (H.ref), immediately after (H0), and 60min (H1) and 120min (H2) after the insult. The same imaging protocol was repeated three times in the Sham group: initial (H.ref), at 60min (H1) and at 120min (H2). For the DWI, seven adjacent axial slices (1mm thickness) were acquired between −5 mm and +1 mm relative to the Bregma, using ultrafast echo planar imaging (EPI; repetition time = 2200msec; echo time = 30msec; field of view = 30×30mm^2^; segments=2; acquisition matrix = 128×128; 16 averages). Parameters of the diffusion sensitization were: duration (δ) = 1.39msec, separation (Δ) = 15msec, b~0 sec/mm^2^ (reference) or b = 800 sec/mm^2^. The DWI acquisition lasted about 8min, after which ADC maps were computed as the mean of the ADCs observed in each of three orthogonal directions.

### Image analysis

Two ROI were manually drawn on the ADC maps: one ROI included the cortex (cortical-ROI) of each hemisphere and one included the whole brain tissue, except ventricular (brain ROI). The two ROIs were drawn on the same single image of the 7 coronal acquisitions. As rats in the TBI group had MRI measurements before and after TBI, images acquired before the trauma were co-registered to T2-weighted images acquired after the trauma using an automated, normalized cross correlation algorithm (co-register function in SPM12 free software, distributed under the terms of the GNU General Public License as published by the Free Software Foundation).

The PRM analysis was conducted as previously described by Galbàn et al. (12). First, we calculated the threshold that designated a significant change in ADC for each rat. To determine this threshold, we performed two linear least squares regression analyses between ADC values obtained at the reference time point (H.ref) and at each of the two following time points H1 and H2 for each Sham-operated rat. The mean threshold and its 95% confidence interval were then determined from these 14 linear regression lines. Second, PRM analysis was applied to each rat in the TBI group using the threshold calculated using sham group. We computed, voxel-wise, ADC differences between the reference time and H0, H1 or H2. If these intra-voxel ADC differences were comprised within the 95% confidence interval of thresholds as defined above, the voxel was considered stable and labeled green. Voxels where ADC differences exceeded that threshold were labeled red. Conversely, voxels where ADC differences were lower than threshold were labeled blue. Results were expressed as the percentage of green, red and blue voxels within each ROI. Therefore, three PRM maps were obtained in the TBI group: between H.ref and H0 (TBI_PRM-H0_), between H.ref and H1 (TBI_PRM-H1_) and between H.ref and H2 (TBI_PRM-H2_). Two PRM maps were obtained in the Sham group: between H.ref and H1 (Sham_PRM-H1_) and between H.ref and H2 (Sham_PRM-H2_).

### Statistical analyses

Data are expressed as mean±SEM. The statistical significance of temporal changes was analyzed using a one-way analysis of variance for repeated measurements (time x groups). Each value was compared to that obtained at the reference time (H.ref) using the Tukey test as a post-hoc test (intragroup analysis). Intergroup analysis was performed using the non-parametric Mann-Whitney test as a post hoc test. (Prism, Graphpad Software Inc., La Jolla, CA, USA).

## Results

The two groups of rats were comparable in terms of their physiological data at H.ref, except for PaO_2_ which remained physiologic or supra physiologic in both group (Table 1).

**Table 1.** Physiological data.

| *Variables \** | *Sham group* | *TBI group* | *p value* |
| --- | --- | --- | --- |
| Weight (g) | 408±78 | 450±58 | 0.16 |
| MABP (mm Hg) | 105±12 | 109±14 | 0.35 |
| EtCo<sub>2</sub> (mm Hg) | 30±6 | 32±4 | 0.56 |
| PH | 7.38±0.1 | 7.39±0.10 | 0.69 |
| PaO<sub>2</sub> (mm Hg) | 103±17 | 157±27 | 0.01 * |
| PaCO<sub>2</sub> (mm Hg) | 39±12 | 31±6 | 0.11 |
| H0- T° (°C) | 36.6±0.9 | 36.4±0.7 | 0.31 |

Table 1. Physiological data were comparable, except for PaO_2_ which remained physiologic in both group.

### Averaged ADC analysis

Neither group showed any significant ADC changes over time in the Cortical- and Brain-ROI (intragroup analysis). However, averaged ADC values in cortical-ROI were significantly reduced at H2 in the TBI group compared to the Sham group: 715±24 μm^2^/sec vs. 781±16 μm^2^/sec, respectively (p=0.04; Fig. 1a). This difference was not found when the brain-ROI was considered: 748±27 μm^2^/sec (TBI group) vs. 801±18 μm^2^/sec (Sham group) (p=0.32) (Fig. 1b). Except for the cortical-ROI at H2, mean ADC values were comparable between the TBI and the Sham group for all ROI and at all time-points.

**Fig. 1.**
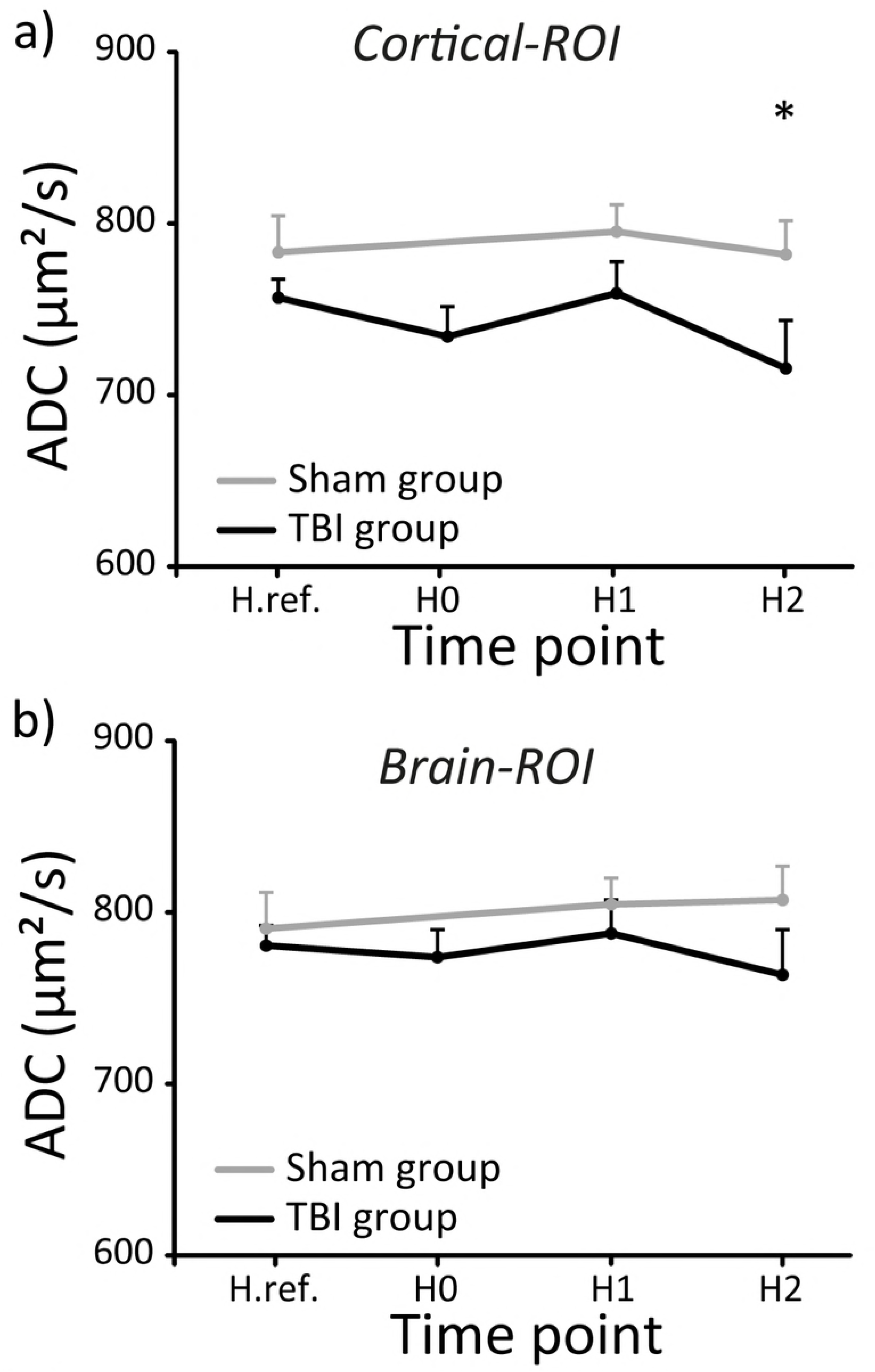
Evolution of mean ADC value. Evolution of mean ADC values in the cortical (a) and brain ROI (b) in TBI and Sham rats. (*: p=0.04, TBI rats vs. Sham rats).

### PRM analysis

In the Sham group, the threshold below which 95% of the voxels are considered stable (PRM0 definition) was 100 μm^2^/sec. Applying PRM in the TBI-group, using this threshold, we found differences between rats in the nature and evolution of post-traumatic edema. Three populations could be distinguished: one population with predominance of voxels reflecting vasogenic edema over the time (red voxels)(Fig.2.a), one population with predominance of voxels reflecting cellular edema over the time (blue voxels)(Fig.2.b), and one population with an initial phase of vasogenic edema followed by a phase of cellular edema (Fig.2.c). In this last population, we could observe a coexistence of vasogenic and cellular edema within the same rat. The TBI group had a higher proportion of voxels with changed ADC values (increased and decreased; respectively red and blue) than Sham group. At H1, differences between TBI_PRM-H1_ and Sham_PRM-H1_ were: red voxels: 16.1±3% vs. 2.7±1% (p=0.005); blue voxels: 11.1±4.7% vs 3±1.8% (p=0.04) (Fig. 3). The total number of voxels found beyond the PRM threshold (i.e. red + blue voxels) were 27.5±4% vs 5.1±1%, respectively (p<0.001). Differences were also revealed at H2 between TBI_PRM-H2_ and Sham_PRM-H2_: red voxels: 9.8%±2.4 vs. 3.5%±2, (p=0.016); blue voxels: 18.4±7.3% vs 1.7±1.6% (p=0.03) (Fig. 3). The total number of voxels beyond the PRM threshold were 29.7±1% vs 5.1±1%, respectively (p<0.001).

**Fig. 2.**
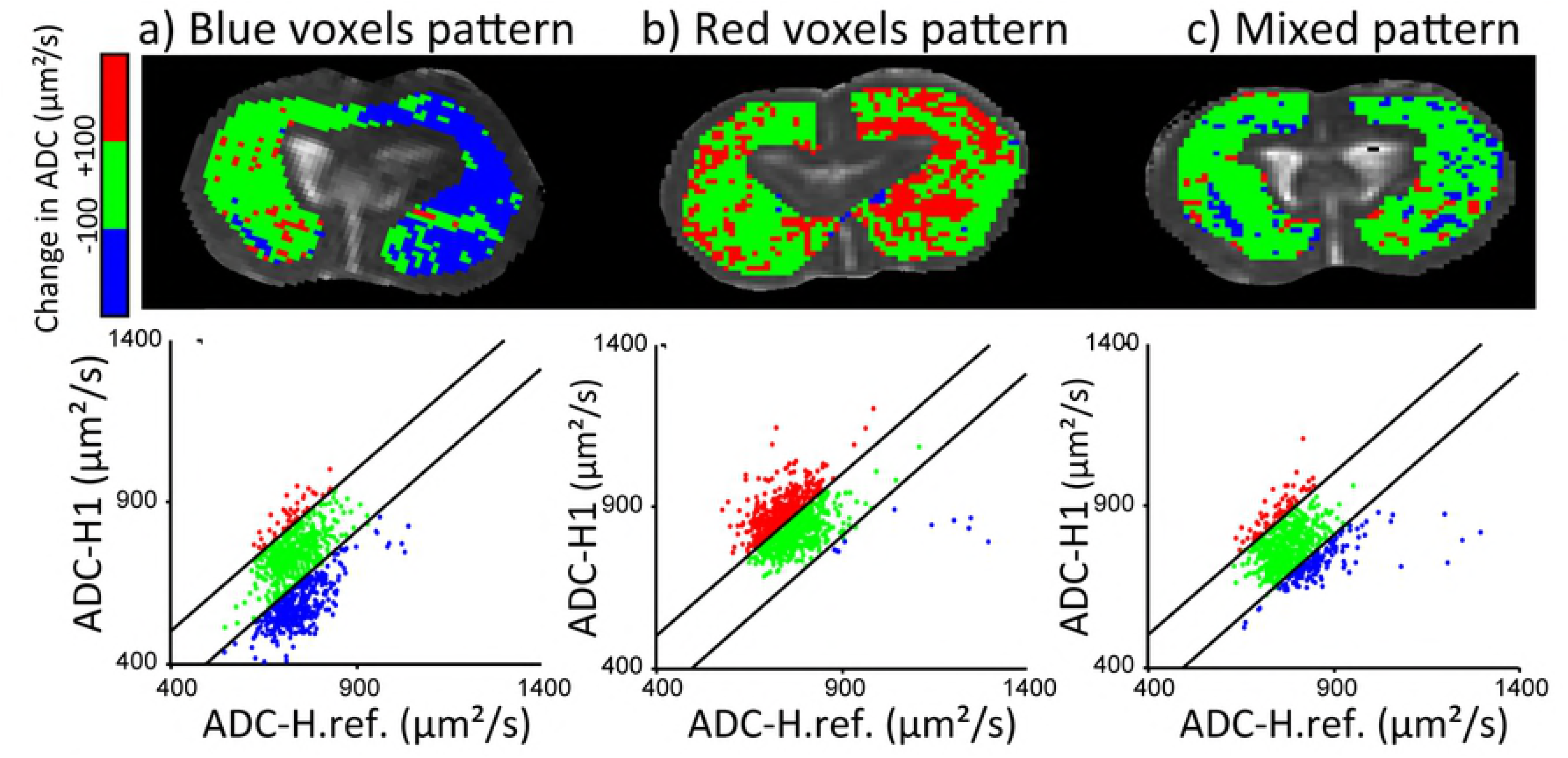
PRM analysis. PRM analysis detects the edematous process and reveals different patterns. Three different rats are represented: a) blue voxel pattern, b) red voxel pattern, c) mixed pattern. The PRM analysis was performed at H1. Displayed images correspond to the PRM color-coded brain-ROI superimposed onto the ADC map. For each image, the representative scatter plot showing the distribution of ADC at H.ref (horizontal axis) compared with that at H1 (vertical axis).

**Fig. 3.**
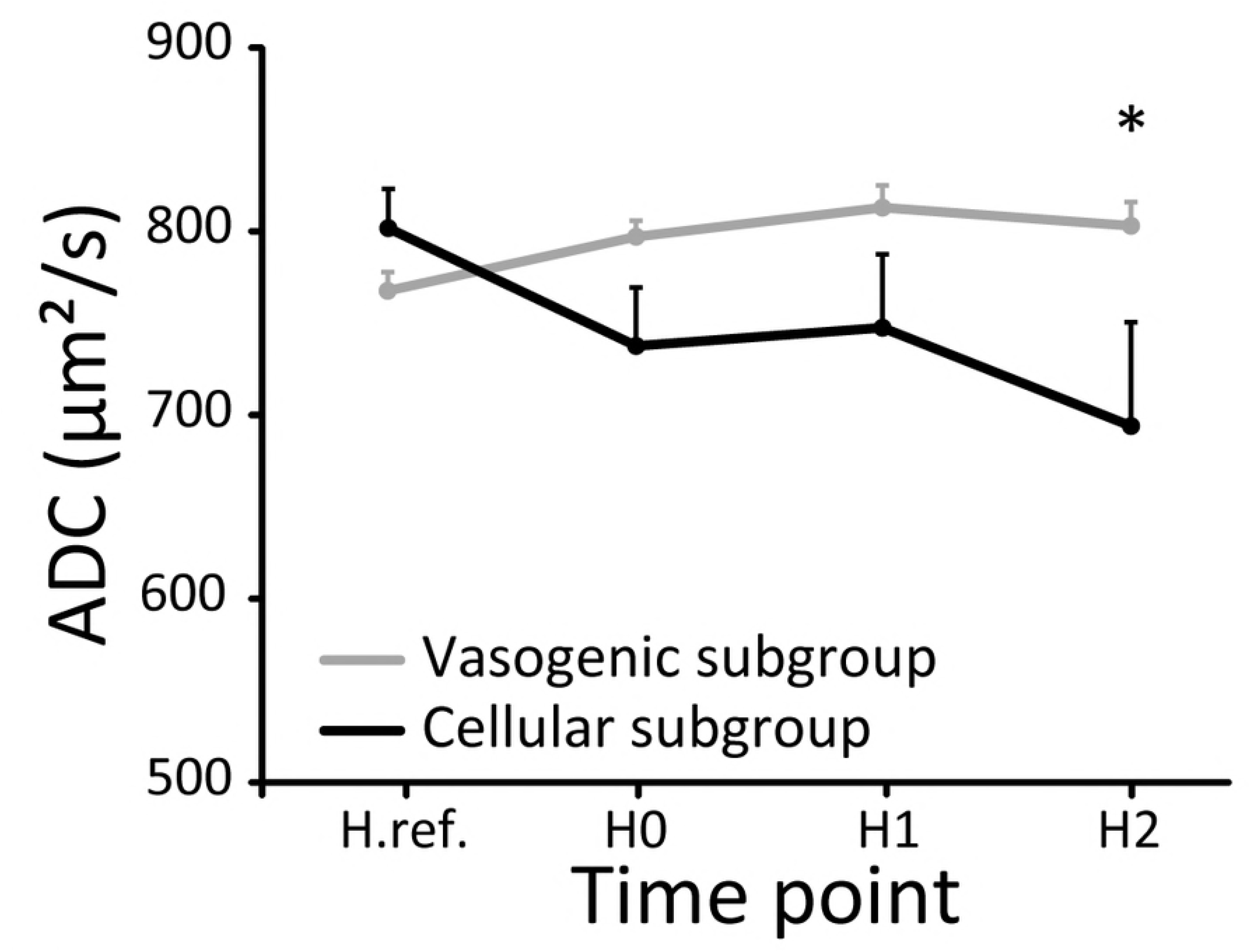
Quantitative representation of the percentage of red and blue voxels in the TBI and Sham groups. The TBI group displayed a sustained higher proportion of red voxels and blue voxels at H1 and H2 compared to the Sham group. Edema in the TBI group is composed of both vasogenic and cellular patterns throughout the time course, though with a peak of red voxels at H1 (*: p<0.05 TBI rats versus Sham rats).

### Subgroup analysis

To further characterize the development of edema in the TBI group, rats in this group were assigned to one of the two following subgroups according to the evolution of their mean ADC value between H.ref and H2 in the brain-ROI: 1) rats presenting an increase in mean ADC were assigned to the vasogenic edema subgroup, and 2) those presenting a decrease in mean ADC to the cellular edema subgroup. Using this approach, 11 rats were assigned to the vasogenic edema subgroup and 7 to the cellular edema subgroup.

As expected, intra-group analysis revealed significant changes in the mean ADC value within both groups over time. In the vasogenic edema subgroup (p=0.001), mean ADC values had increased at H0, H1 and H2 compared to H.ref (H.0 vs H.ref: mean difference= 31.1 μm^2^/sec, q=3.9, 95%CI: [61.1,1.26] μm^2^/sec; H1 vs H.ref: mean difference= 45.7 μm^2^/sec, q=5.8, 95%CI: [75.7, 15.8] μm^2^/sec; and H2 vs. H.ref: mean difference= 35.7 μm^2^/sec, q=4.5, 95%CI: [65.7, −5.8] μm^2^/sec). In the cellular edema subgroup (p=0.001), mean ADC values had decreased only at H2 compared to H.ref (H.2 vs H.ref: mean difference= 140.3 μm^2^/sec, q=5.9, 95%CI: [45.8, 234.8] μm^2^/sec). The evolution over time of the mean ADC values therefore differed between the two subgroups, in line with the subgroup definition, (p<0.01; Fig. 4), with a significantly lower mean ADC at H2 in the cellular edema subgroup than in the vasogenic edema subgroup (vasogenic subgroup: 802.7±10 μm^2^/sec, 95%CI: [780.4, 825.1] μm^2^/sec, cellular subgroup: 694.3±55 μm^2^/sec, 95%CI: [551.9, 836.6] μm^2^/sec, p<0.01; Fig. 4). This difference in ADC values between the subgroups was not found at H1 and H0.

**Fig. 4.**
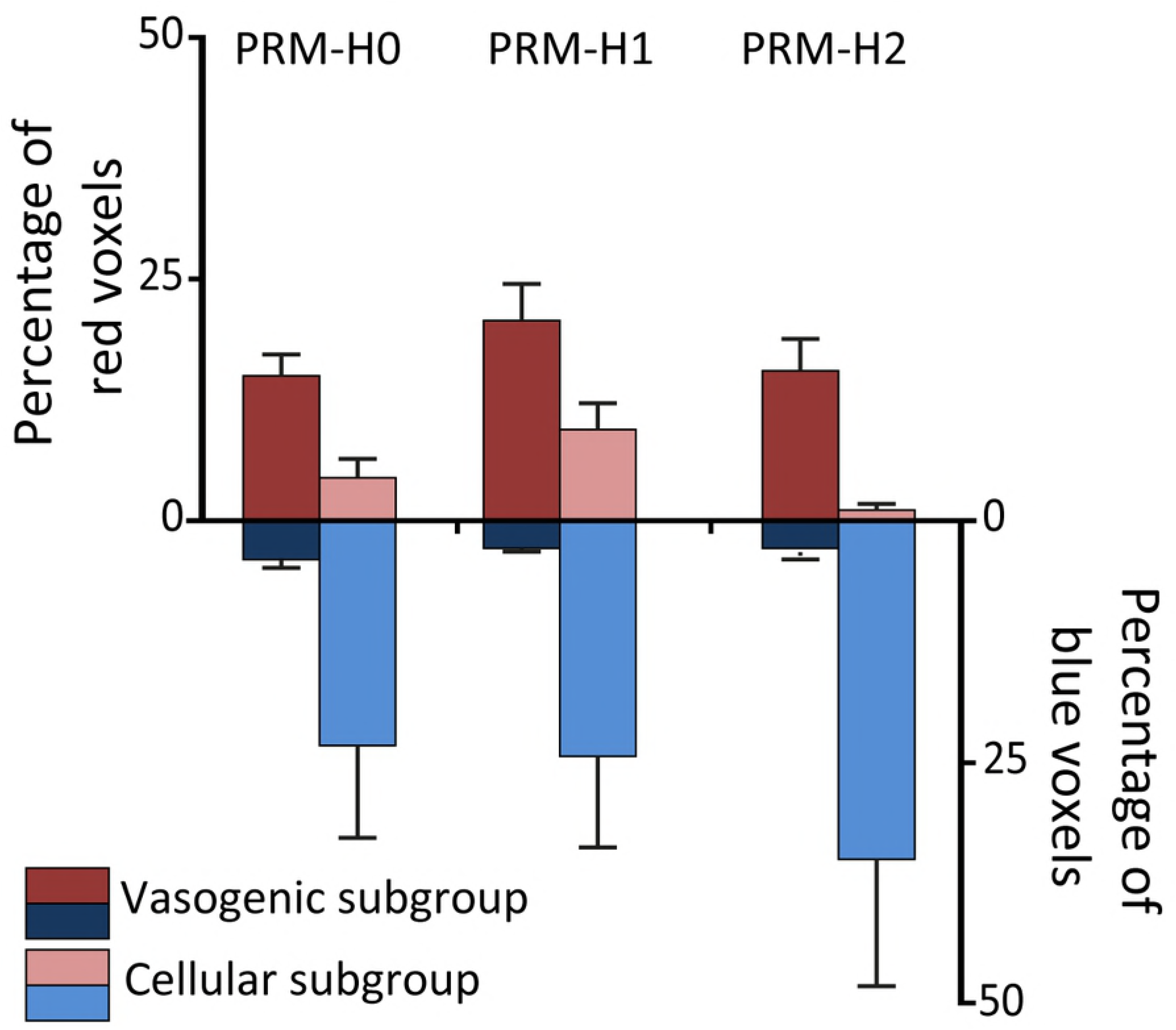
Evolution of mean ADC values within brain-ROI in the cellular subgroup (black) and vasogenic subgroup (grey). Mean ADC values were significantly lower in the cellular subgroup at H2 compared to in the vasogenic subgroup (*: p<0.01).

We then performed PRM analysis on these two subgroups to describe the proportion of each type of edema within each subgroup. The vasogenic edema subgroup presented less than 5% of blue voxels and a proportion of red voxels above 15% throughout the experiment. Conversely, the cellular edema subgroup presented a sustained proportion of more than 25% of blue voxels with less than 8% of red voxels (Fig. 5). The evolution in percentages of blue and red voxels also significantly differed between the two subgroups (p<0.01 and p<0.01; Fig. 5). While the vasogenic subgroup presented a higher percentage of red voxels compared to the cellular subgroup at H0 and H2 (14.7±2 % vs 4.3±2 % at H0; 15.4±3 % vs 1.5±1 % at H2), the cellular subgroup presented a higher percentage of blue voxels compared to the vasogenic subgroup at H1 and H2 (24.2±10 % vs 2.4±1 % at H1; 34.7±14 % vs 2.8±0.4 % at H2). Altogether, after TBI, rats exhibit a mixed pattern of red and blue voxels, with a clear predominance of one or the other.

**Fig. 5.**
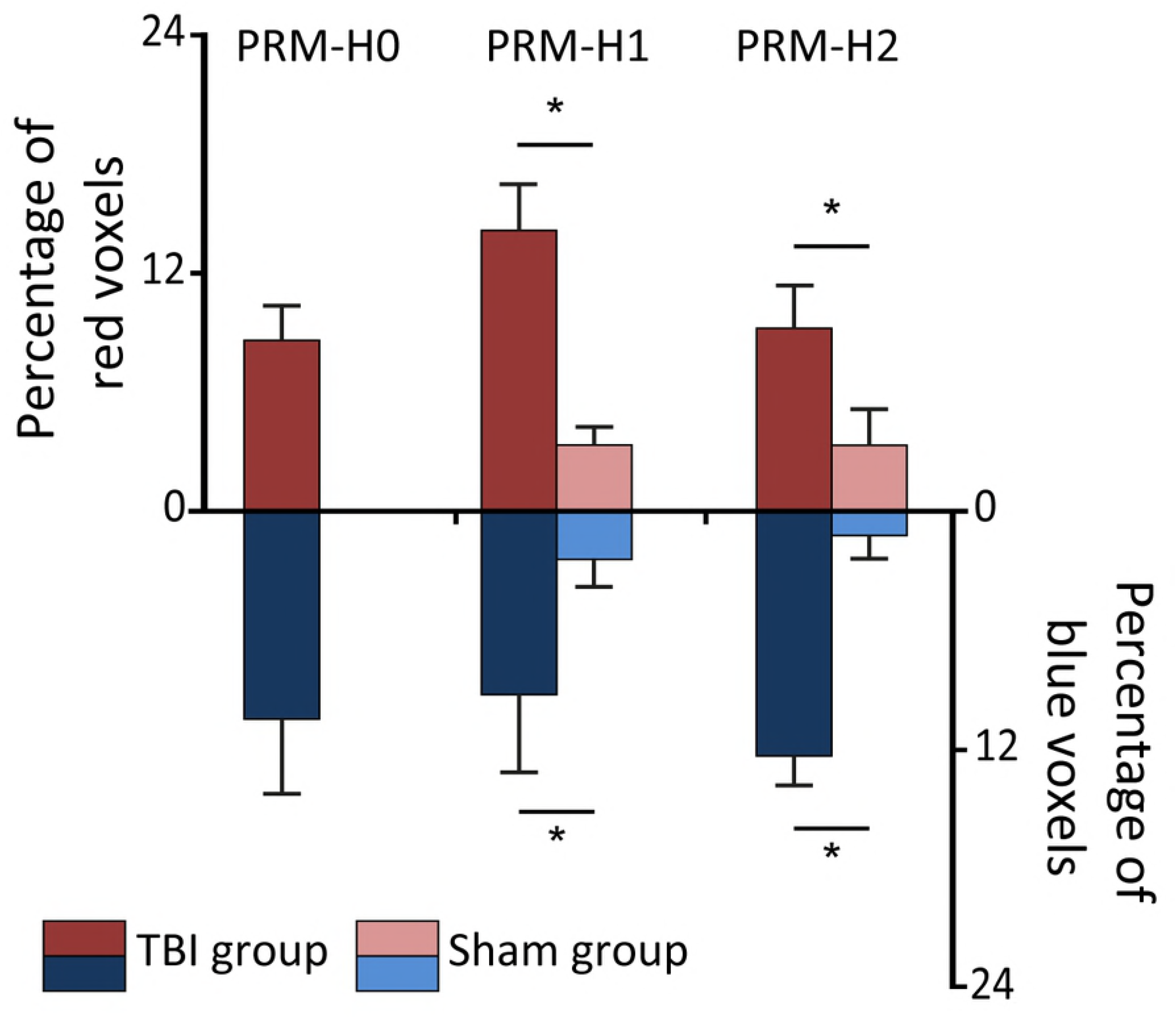
Quantitative representation of the percentage of red and blue voxels in the vasogenic subgroup and cellular subgroup. The vasogenic subgroup displayed a higher proportion of red voxels compared to the cellular subgroup at H0 and H2. The cellular subgroup sustained a higher proportion of blue voxels compared to the vasogenic subgroup at H1 and H2 (*: p<0.05 TBI rats versus Sham rats). Both subgroups displayed a peak of red voxels at H1.

## Discussion

Averaging ADC values in the whole brain failed to detect the occurrence of brain edema in our rat model of diffuse TBI. A voxel-wise basis approach (PRM) could however detect the nature of post-traumatic brain edema, i.e., vasogenic, cellular or a mixed pattern, and is able to provide temporal and spatial information concerning evolution of this edema. These findings indicate that PRM could provide detailed and accurate information of the nature of brain edema after trauma and its evolution. This may be of value in clinical practice to individualize the management of brain-injured patients.

Impact acceleration model is known to induce diffuse traumatic brain injury (13, 14) with diffuse axonal injuries (15) without preliminary skull opening. This model is used for almost twenty years as it is close to what could be observed in a clinical point of view. Most previously published studies have focused on mean ADC values to assess cerebral edema and limited their investigations to specific small ROIs such as cortex and striatum (3, 6, 7). Using a cortical-ROI, our findings are consistent with previous studies showing a lower value of mean ADC in the TBI group at H2 compared to Sham-operated rats (7). However, when using this classical mean-value approach on a larger ROI, we found no difference between the TBI and the Sham groups, suggesting an absence of cerebral edema in the TBI groups. These results suggest that the mean-value approach can be used to measure change in ADC only in very specific areas. In addition, ADC values in the cortex at H2 exhibited high standard deviations, suggesting an inter-subject heterogeneity in the development of edema, as previously found (3, 7). Conversely, the PRM analysis was able to describe the development of cerebral edema in a large ROI after diffuse TBI. It also showed that these ROIs contained areas of heterogeneously evolving ADC, suggesting that mean ADC approach, which averages out positive and negative ADC changes, may have led to misleading conclusions. Moreover, the mean-value approach is limited to one quantitative estimate (the mean ADC in this case) whereas PRM also allows the measurement of the proportion of voxels which vary above a predefined ADC threshold.

We were surprised to find almost no difference between the TBI and the Sham-operated rats in terms of mean ADC values across a whole brain ROI. The results of the PRM analysis showed a spatial heterogeneity and may explain this absence of any difference. Only 15% of voxels exhibited ADC changes during the experiment in the TBI group at H2. With a mean-value approach across the entire ROI, significant modifications of ADC in a small proportion of voxels would be hidden by a high proportion of stable voxels (partial volume effect). Our results also show that a mixed pattern of vasogenic and cellular edemas is generally observed which would further reduce the ability of a mean ADC estimate to detect the presence of edema. Moreover, our results suggest that post traumatic edema is not restricted to specific brain areas usually chosen for ROI, such as the cortex or striatum, but could be spread throughout the whole brain (Fig.2). The spatial and temporal evolution of edema could thus be estimate by PRM within every animal. The focus on specific ROI could therefore lead to misinterpretations, eluding the complexity of post-traumatic cerebral edema.

One important finding of our study is the presence of red voxels in almost every rat. This elevation of ADC could be transient (Fig.2.c), according to previous studies (3), or could be sustained across the experiment. DWI in diffuse brain injury reported initial predominant cellular edema with decreased ADC values (3, 5–7). Recently Hudak et al. showed that both cellular and vasogenic edema could coexist in the first week after severe TBI. Interestingly, the vasogenic edema was prominent in these patients (8). In this work, the only parameter that influenced the development of cerebral edema was the severity of the TBI, as measured by the initial Glasgow Score Scale. In our study, all rats in the TBI group were comparable and subjected to the same TBI protocol yet the proportion of vasogenic or cellular subpopulations evolved. Moreover, physiological data showed no differences between the two subgroups, with the exception of PaO2 which remains physiologic in both groups (Table 1) and could not explain the difference in the evolution of the cerebral edema. This finding leads to suspect an impact of individual susceptibility for developing specific edematous response after TBI. Nevertheless, a PRM analysis of early ADC data was useful in estimating the edematous pattern two hours after diffuse traumatic brain injury. This finding could be relevant with regards early classification according to edematous status in preclinical studies. The use of more homogeneous groups could certainly improve the evaluation of neuroprotective strategies.

From the clinical standpoint, distinction between vasogenic and cellular edema could be the next key issue for the management of brain injury: recognition of edema type could lead to specific therapies such as lowering blood pressure or reducing fluid administration in the event of vasogenic edema (16). Moreover, recent studies highlighted the potential beneficial effect of specific molecules, such as inhibitors of aquaporins, on water accumulation after cellular edema (17–19). As demonstrated in 2012 by Galbán et al., PRM analysis is not restricted to MRI use and can be successfully performed on whole-lung computed tomography (CT) scans, to differentiate between specific chronic obstructive pulmonary disease phenotypes (20). In clinical practice, MRI is not always accessible for the severely injured patient and CT scan remains the routine examination. Recently, Hakseung et al. (23) was able to quantify brain tumor edema using CT scans on a pediatric TBI population. These studies demonstrate the feasibility of using PRM analysis on CT scan in order to monitor brain edema of TBI patients. Evaluation of PRM on CT scans in neurocritical care has never been performed yet could be helpful for the clinician. Indeed, PRM analysis could help physicians to manage patients with severe TBI depending on the mechanisms involved in the brain edema formation. The question of the reference data for TBI patients remains unsolved at the moment, but some strategies are tested to progress in this way and particularly the use of reference atlas of the human brain, or the use of the symmetry with the non-injured part of the patients brain. Another interesting application for PRM is as an imaging biomarker to test the efficiency of therapies, as previously shown in oncology (21, 22). Prognostication of functional outcome in patients suffering from non-traumatic intracerebral hemorrhage could also be relevant for physicians (23).

In conclusion, our findings show that monitoring post-traumatic cerebral edema by mean ADC values in specific ROIs could be misleading. PRM in a voxel-based approach appears more accurate than the classical averaging approach to detect both vasogenic and cellular edema. Our PRM analysis further highlighted that the impact-acceleration model induces several edematous patterns and suggested an individual specificity with regards trauma-related cerebral edema. Utilization of PRM may help physicians and researchers to early sort cerebral edema in homogeneous subgroups.

## List of abbreviations used in the text

ADC: apparent diffusion coefficient
CE: cerebral edema
ICP: intracranial pressure
MRI: magnetic resonance imaging
PRM: parametric response map
ROI: region of interest
SEM: standard error of the mean
TBI: traumatic brain injury

## References

1. Corrigan JD, Selassie AW, Orman JAL. The epidemiology of traumatic brain injury. J Head Trauma Rehabil. avr 2010;25(2):72–80.

2. Klatzo I. Presidental address. Neuropathological aspects of brain edema. J Neuropathol Exp Neurol. janv 1967;26(1):1–14.

3. Barzó P, Marmarou A, Fatouros P, Hayasaki K, Corwin F. Contribution of vasogenic and cellular edema to traumatic brain swelling measured by diffusion-weighted imaging. J Neurosurg. déc 1997;87(6):900–7.

4. Ito J, Marmarou A, Barzó P, Fatouros P, Corwin F. Characterization of edema by diffusion-weighted imaging in experimental traumatic brain injury. Journal of Neurosurgery. 1 janv 1996;84(1):97–103.

5. Albensi BC, Knoblach SM, Chew BG, O’Reilly MP, Faden AI, Pekar JJ. Diffusion and high resolution MRI of traumatic brain injury in rats: time course and correlation with histology. Exp Neurol. mars 2000;162(1):61–72.

6. Verdonck O, Lahrech H, Francony G, Carle O, Farion R, Van de Looij Y, et al. Erythropoietin protects from post-traumatic edema in the rat brain. J Cereb Blood Flow Metab. juill 2007;27(7):1369–76.

7. Bouzat P, Francony G, Thomas S, Valable S, Mauconduit F, Fevre M-C, et al. Reduced brain edema and functional deficits after treatment of diffuse traumatic brain injury by carbamylated erythropoietin derivative. Crit Care Med. sept 2011;39(9):2099–105.

8. Hudak AM, Peng L, Marquez de la Plata C, Thottakara J, Moore C, Harper C, et al. Cytotoxic and vasogenic cerebral oedema in traumatic brain injury: Assessment with FLAIR and DWI imaging. Brain Inj. 24 juill 2014;28(12): 1602–9.

9. Unterberg AW, Stover J, Kress B, Kiening KL. Edema and brain trauma. Neuroscience. 2004;129(4):1019–27.

10. Dixon CE, Clifton GL, Lighthall JW, Yaghmai AA, Hayes RL. A controlled cortical impact model of traumatic brain injury in the rat. J Neurosci Methods. oct 1991;39(3):253–62.

11. Long JA, Watts LT, Chemello J, Huang S, Shen Q, Duong TQ. Multiparametric and longitudinal MRI characterization of mild traumatic brain injury in rats. J Neurotrauma. 15 avr 2015;32(8):598–607.

12. Li W, Watts L, Long J, Zhou W, Shen Q, Jiang Z, et al. Spatiotemporal changes in blood-brain barrier permeability, cerebral blood flow, T2 and diffusion following mild traumatic brain injury. Brain Res. 01 2016;1646:53–61.

13. Moffat BA, Chenevert TL, Lawrence TS, Meyer CR, Johnson TD, Dong Q, et al. Functional diffusion map: a noninvasive MRI biomarker for early stratification of clinical brain tumor response. Proc Natl Acad Sci USA. 12 avr 2005;102(15):5524–9.

14. Galban CJ, Chenevert TL, Meyer CR, Tsien C, Lawrence TS, Hamstra DA, et al. The parametric response map is an imaging biomarker for early cancer treatment outcome. Nat Med. mai 2009;15(5):572–6.

15. Marmarou A, Foda MA, van den Brink W, Campbell J, Kita H, Demetriadou K. A new model of diffuse brain injury in rats. Part I: Pathophysiology and biomechanics. J Neurosurg. févr 1994;80(2):291–300.

16. Foda MA, Marmarou A. A new model of diffuse brain injury in rats. Part II: Morphological characterization. J Neurosurg. févr 1994;80(2):301–13.

17. Meythaler JM, Peduzzi JD, Eleftheriou E, Novack TA. Current concepts: diffuse axonal injury-associated traumatic brain injury. Arch Phys Med Rehabil. oct 2001;82(10):1461–71.

18. Jungner M, Grände P-O, Mattiasson G, Bentzer P. Effects on brain edema of crystalloid and albumin fluid resuscitation after brain trauma and hemorrhage in the rat. Anesthesiology. mai 2010;112(5):1194–203.

19. Manley GT, Fujimura M, Ma T, Noshita N, Filiz F, Bollen AW, et al. Aquaporin-4 deletion in mice reduces brain edema after acute water intoxication and ischemic stroke. Nat Med. févr 2000;6(2):159–63.

20. Papadopoulos MC, Manley GT, Krishna S, Verkman AS. Aquaporin-4 facilitates reabsorption of excess fluid in vasogenic brain edema. FASEB J. août 2004; 18(11):1291–3.

21. Badaut J, Fukuda AM, Jullienne A, Petry KG. Aquaporin and brain diseases. Biochim Biophys Acta. mai 2014;1840(5):1554–65.

22. Galbán CJ, Han MK, Boes JL, Chughtai KA, Meyer CR, Johnson TD, et al. Computed tomography-based biomarker provides unique signature for diagnosis of COPD phenotypes and disease progression. Nat Med. nov 2012; 18(11):1711–5.

23. Kim H, Kim G, Yoon BC, Kim K, Kim B-J, Choi YH, et al. Quantitative analysis of computed tomography images and early detection of cerebral edema for pediatric traumatic brain injury patients: retrospective study. BMC Med. 22 oct 2014;12:186.

24. Lemasson B, Galbán CJ, Boes JL, Li Y, Zhu Y, Heist KA, et al. Diffusion-Weighted MRI as a Biomarker of Tumor Radiation Treatment Response Heterogeneity: A Comparative Study of Whole-Volume Histogram Analysis versus Voxel-Based Functional Diffusion Map Analysis. Transl Oncol. 2013;6(5):554–61.

25. Boes JL, Hoff BA, Hylton N, Pickles MD, Turnbull LW, Schott AF, et al. Image registration for quantitative parametric response mapping of cancer treatment response. Transl Oncol. févr 2014;7(1):101–10.

26. Tsai Y-H, Hsu L-M, Weng H-H, Lee M-H, Yang J-T, Lin C-P. Functional diffusion map as an imaging predictor of functional outcome in patients with primary intracerebral haemorrhage. Br J Radiol. janv 2013;86(1021):20110644.

